# CytoPb: Segmentation-free and threshold-free analysis of highly multiplexed images of biological tissue

**DOI:** 10.1101/2023.07.26.550516

**Authors:** Paul R Barber, Luigi Dolcetti, Chen-An Yeh, Tony Ng

## Abstract

We present a novel method and code, CytoPb, for the analysis of multiplexed images that is segmentation-free and threshold-free and produces results that correlate well with existing methods but is faster to run, needs minimal training and removes subjectivity associated with the validation of single-cell segmentation and thresholding. In three data sets cell abundance measures correlate with previous methods (Pearson’s coef. >0.75). Neighbourhood analysis and cell-cell proximity are possible.

## Main Text

The field of multiplexed imaging encompasses many rapidly evolving techniques, such as spatial transcriptomics and imaging mass cytometry (IMC) and is enabling the visualization and quantitative measurement of multiple molecular and cellular features within a single sample. The quantification of multiple key features of the tissue, such as cell proliferation, apoptosis, angiogenesis, and immune cell infiltration is crucial for understanding the underlying mechanisms of disease such as cancer. This approach provides a comprehensive view of the complex biological processes and helps gain deeper insights into the mechanisms of cancer development, progression, and response to therapy.

Multiplexed imaging has become an indispensable tool in cancer research, allowing for the simultaneous analysis of multiple biomarkers. However, current methods face analysis challenges that involve single-cell segmentation and channel thresholding (gating), which sometimes force subjective decisions and reduced analysis reproducibility. In this paper, we present a new analysis method, called CytoPb, that overcomes these challenges by using a segmentation-free and threshold-free method based on marker and cell-type probability maps, and allows the usual follow-on processing of tissue neighbourhoods and cell-cell proximity and interactions.

Previous method for cell-type determination segment images into single-cell regions using a nuclear and/or membrane marker, and often use a random forest or deep learning classifier^1,2 3^ that may exploit many image channels. The success of this process can be subjective and lacks a gold standard such that the user is left to judge segmentation quality by visual inspection. Thresholding then determines if the segmented cells are positive or negative for particular markers (e.g. cluster of differentiation (CD) markers). This process is also somewhat subjective with the user left to visually inspect the markers within cell regions to decide the cut-off between positive and negative. These decisions can be difficult if the marker staining does not clearly mark cell nuclei or boundaries, and positivity is not just a function of mean region intensity but also of spatial distribution. Making difficult and irreversible decisions early in the analysis pipeline is not desirable. The Toponomics method^4^ exploited pixel-based analysis rather than segmentation but did use thresholds on a pixel-by-pixel basis.

It is advantageous to use a probabilistic or fuzzy logic approach to evade or delay decisions until the last possible stage of the analysis. Here, that approach is used to produce the usual cell abundance measurements that agree well with previous methods (correlation coefficient >0.75), cell type maps as well as neighbourhood analysis and cell proximity/interaction measurement. This method copes with all the complex marker patterns usually used in immunology, including enforcing that a cell type should be negative for certain markers, in a cell-type marker matrix. Neighbourhood analysis is achieved by clustering image regions according to cell type content. Proximity analysis can be performed by generating scale space overlap signatures^5^.

The method also offers significant time saving, running completely automatically in a fraction of the time taken to train a segmentation classifier and perform manual channel thresholding. It can also automatically compensate for certain batch effects between imaging runs (e.g. staining efficiency) therefore minimising manual intervention. Standard methods do not scale well if there are any variations in batches including changes to the marker panel. The current implementation runs purely in R with open-source packages and the current run speed is 1-2 minutes per image with a 4 GHz processor and >8 GB of RAM, and so is easily run on a current desktop PC. A speed-up of several factors could be achieved on multi-core computers or clusters with multi-threaded code.

Figure 1 shows a schematic overview of the method by which the raw data from marker channels (the metallic tag channels of IMC or fluorescence channels or any image of marker distribution) become marker probability maps through the estimation of the marker levels that represent definite negative and positive expression through standard image processing operations that identify consolidated areas of high levels of expression and filter out random and single pixel noise. These levels are, in general, much easier to determine than a threshold that divides positive from negative and can be manually adjusted. The resolution of the images may be slightly reduced with Gaussian filtering to account for the size of biological cells that are much larger than single pixels in this kind of imaging.

**Figure 1:**
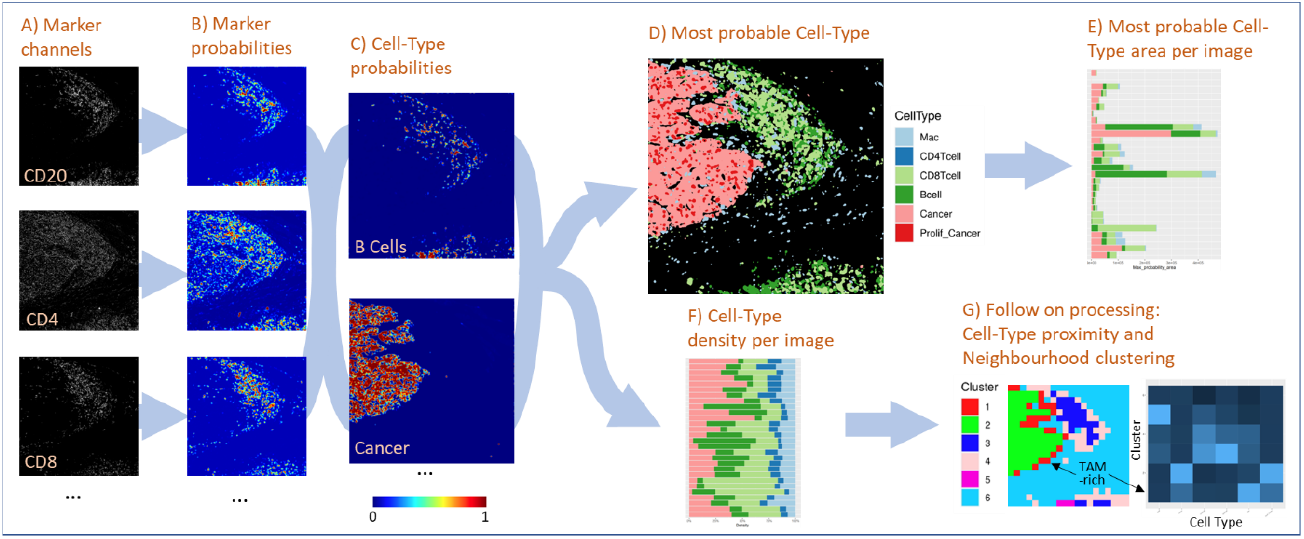
CytoPb workflow from Marker channel intensities (A) to fuzzy marker probabilities (B) that are combined through AND and OR fuzzy logical operations to produce cell-type probability maps (C). Maps of the most probable cell type per pixel can be produced for visualisation (D), which can also be quantified for many images in an experiment (E). Alternatively, all probability information can be retained and totalled to produce cell type density (abundance) per image (F) and further processing can be performed to calculate characteristic proximities between cell types and neighbourhood clustering (G). Highlighted is Cluster 1 which contains areas of tumour associated macrophages (TAM) on the periphery on the cancer cell mass, as confirmed in the cell-type per cluster heatmap.

These marker probability maps are combined by fuzzy logic operations to produce maps of cell type probability as defined by a cell-type matrix. It is trivial then to determine the most likely cell type for any given pixel by choosing the cell type with highest probability (greater than some minimum, say 0.1, to avoid unlikely types). Summing the probability per cell type gives an estimate of cell abundance. The cell type maps produced can also be used for further processing such as region analysis by clustering to identify tissue types^6^.

Enhancements could incorporate custom spatial filters (other than Gaussian) and intensity filters (other than sigmoidal) to account for markers that are described as low/med/high rather than negative/positive, or to supress bright artefacts such as autofluorescence.

In Figure 2 the results from this method are compared with three data sets with different acquisition and segmentation methods: Bone marrow IMC using the Steinbock analysis package<u>^2^</u>, Head and Neck Cancer tissue using RUNIMC^7^, and the Colorectal Cancer tissue CODEX data and method of Schurch et al.^8^ Efforts were made to match the cell-type gating strategies. The segmentation methods included corrections for batch effects between samples, CytoPb use no corrections.

**Figure 2:**
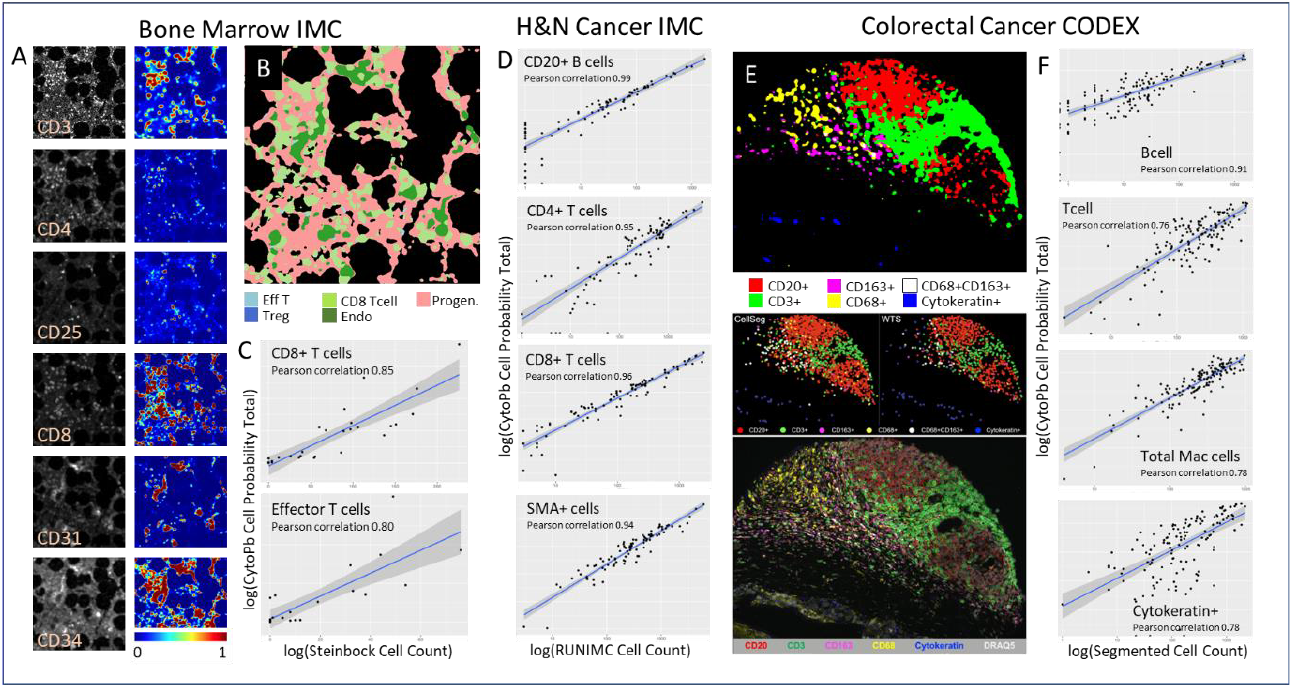
Demonstration of CytoPb on three datasets and correlations with traditional segmented and thresholded cells. Example Imaging CyTof (IMC) data of bone marrow and fuzzy channel maps (A). Example cell type map (B). Correlations with segmentation method using the Steinbock pipeline from 29 images (C). Correlations with the RUNIMC segmentation method from 90 images from head and neck cancer IMC (D). CytoPB produced-cell type map from CODEX data (E top) and comparison with previously published result (from Lee et al. BMC Bioinformatics, 2022, under the terms of the Creative Commons Licence) using the CellSeg algorithm (E middle left) and watershed and thresholding (WTS, E middle right) and marker intensities image (E bottom). Correlations with segmentation method from 140 CODEX images (Schurch et al. Cell, 2020, F). “Total Mac cells” includes a sum of all cells in the CD163+, CD68+ and CD68+CD163+ groups.

In conclusion, we present an alternative method for the analysis of multiplexed images of tissue that correlates well with single cell segmentation but importantly allows the avoidance of sometimes subjective segmentation and thresholding during the analysis. For this reason, it is expected to be more reproducible. The CytoPb source code is available here: https://github.com/paulbarber/CytoPb.

## Methods (Supplementary Info, with section headers)

All processing is performed in R using the packages mentioned below.

### Data Import

Data are imported from their native format and converted into multi-channel tiff images if necessary. For IMC data, the imctools package was used.

### Marker Levels (Channel Ranges)

The estimation of the marker levels that represent definite negative and positive expression is automatic by the approximate segmentation of some definite foreground area. This involves median filtering and normalisation, thresholding and joining 4-pixel connected regions, and removing small regions (<10 pixels). This process is very strict (highly specific, low sensitivity) to find definite positive areas. The positive level is set at the 5% quantile of the selected pixels. The negative value is taken from the mean of the remaining pixels.

### Marker Probability Maps

Probability maps for each channel are produced and are strictly in the range [0,1] using a sigmoid function based on the determined marker levels and a slope parameter. Images are exported for visual inspection. Marker levels can be manually modified if necessary, and new maps generated.

### Cell-Type Probability Maps

Cell types are defined in a csv file by a matrix of +1 or -1 that indicates which markers should be positive and negative for each cell type. Cell-type probability maps are calculated by using fuzzy logic operations (e.g. the min operator performs a logical ‘AND’) on combinations of marker probability maps (or 1 minus the map for a negative marker). The resultant cell maps are also in the range [0,1].

Unsupervised analysis can be performed by pixel clustering (or using regions of mean marker intensity) to aid the selection of cell types.

### Cell Type Plots and Heatmaps

The cell-type membership of each pixel remains ‘fuzzy’ and probabilistic. For cell-type maps the most likely cell type per pixel is chosen by the most likely cell type per pixel. To measure cell abundance (number), the total cell probability (sum over whole image) is used. Cell density can be calculated with respect to the image size or the total tissue probability (determined from the logical OR of all channels, proportional to tissue area).

The outputs from this stage are cell type images (coloured by cell type), data and plots of cell abundance, density, and area per image, and heatmaps of marker intensity per cell type to check for consistency with expectations.

### Neighbourhood clustering

Cell neighbourhoods can be identified by measuring the cell abundances within image regions. This has been performed by forming a grid of image squares (50×50 um) and clustering the cell abundances using kmeans or hierarchical clustering. Heatmaps of cell type content per region can be used to identify neighbourhood types, and maps and plots of neighbourhood type per image are produced.

### Cell-Cell proximity

Measurements of cell-cell proximity can be made on the assumption that the cell-type probability map represents the kernel density estimate (KDE) and calculating the overlap between cell types. By calculating the KDE at different spatial scales, a scale-space overlap signature can be determined that is characteristic of the tendency for two cell types to be collocated. This is performed in CytoPb using Gaussian kernels and the overlap calculated from the pixel-by-pixel minimum of the two images. When moving from a fine scale to a coarser scale (over a sufficient range) the overlap will transition from zero to one (this signal is the “scale signature”). The characteristic scale of the overlap is taken as the maximum gradient of the scale signature.

## Data availability

Colorectal Cancer tissue CODEX data and method of Schurch et al. is available from: https://data.mendeley.com/datasets/mpjzbtfgfr/1 Other data is available from the corresponding author at reasonable request.

## Code availability

The code for the algorithms presented in this paper are available here: https://github.com/paulbarber/CytoPb

## Acknowledgements

We would like to thank Cancer Research UK for funding (CRUK City of London Centre). We would like to thank Shahram Kordasti for the use of bone marrow data.

